# Archaic adaptive introgression in *TBX15/WARS2*

**DOI:** 10.1101/033928

**Authors:** Fernando Racimo, David Gokhman, Matteo Fumagalli, Amy Ko, Torben Hansen, Ida Moltke, Anders Albrechtsen, Liran Carmel, Emilia Huerta-Sánchez, Rasmus Nielsen

## Abstract

A recent study conducted the first genome-wide scan for selection in Inuit from Greenland using SNP chip data. Here, we report that selection in the region with the second most extreme signal of positive selection in Greenlandic Inuit favored a deeply divergent haplotype that is closely related to the sequence in the Denisovan genome, and was likely introgressed from an archaic population. The region contains two genes, *WARS2* and *TBX15*, and has previously been associated with adipose tissue differentiation and body-fat distribution in humans. We show that the adaptively introgressed allele has been under selection in a much larger geographic region than just Greenland. Furthermore, it is associated with changes in expression of *WARS2* and *TBX15* in multiple tissues including the adrenal gland and subcutaneous adipose tissue, and with regional DNA methylation changes in *TBX15*.

## Introduction

To identify genes responsible for biological adaptations to life in the Arctic, Fumagalli *et al*. (Fumagalli, et al. 2015) scanned the genomes of Greenlandic Inuit (GI) using the population branch statistic (PBS)(Yi, et al. 2010), which detects loci that are highly differentiated from other populations. Using this method, they found two regions with a strong signal of selection. One region contains the cluster of *FADS* genes, involved in the metabolism of unsaturated fatty acids. Several of the SNPs with the highest PBS values in this region were shown to be significantly associated with different phenotypes including fatty acid profiles, weight and height.

The other region contains *WARS2* and *TBX15*, located on chromosome 1. *WARS2* encodes the mitochondrial tryptophanyl-tRNA synthetase. *TBX15* is a transcription factor from the T-box family and is a highly pleotropic gene expressed in multiple tissues at different stages of development. It is required for skeletal development (Singh, et al. 2005) and deletions in this gene cause Cousin syndrome, whose symptoms include craniofacial dysmorphism and short stature (Lausch, et al. 2008). *TBX15* also plays a role in the differentiation of brown and brite adipocytes (Gburcik, et al. 2012). Brown and brite adipocytes produce heat via lipid oxidation when stimulated by cold temperatures, making *TBX15* a strong candidate gene for adaptation to life in the Arctic. SNPs in or near both of these genes have also been associated with numerous phenotypes in GWAS studies – in particular waist-hip ratio and fat distribution in Europeans (Heid, et al. 2010) and ear morphology in Latin Americans (Adhikari, et al. 2015).

Multiple studies have shown extensive introgression of DNA from Neanderthals and Denisovans into modern humans (Green, et al. 2010; Meyer, et al. 2012; Prüfer, et al. 2014; Sankararaman, et al. 2014; Vernot and Akey 2014; Racimo, et al. 2015). Many of the introgressed tracts have been shown to be of functional importance and may possibly be examples of adaptive introgression into humans, including genes involved in immunity (Abi-Rached, et al. 2011; Dannemann, et al. 2015), genes associated with skin pigmentation (Sankararaman, et al. 2014; Vernot and Akey 2014), and *EPAS1*, associated with high-latitude adaptation in Tibetans (Huerta-Sánchez, et al. 2014). It has been hypothesized that Archaic humans were adapted to cold temperatures (Steegmann, et al. 2002). Therefore, in this paper we examine if any of the selected genes in Inuit (Fumagalli, et al. 2015) may have been introduced into the modern human gene pool via admixture from archaic humans, i.e. Neanderthals or Denisovans (Meyer, et al. 2012; Prüfer, et al. 2014). We will show that the *WARS2/TBX15* haplotype that is at high frequency in GI was likely introgressed from an archaic human population. We will also show that the selection affecting this haplotype is relatively old, resulting in a high allele frequency in other New World populations and intermediate allele frequencies in East Asia. Finally, functional genomic analyses suggest that the selected archaic haplotype may affect the regulation of expression of *TBX15* and *WARS2*, and is associated with phenotypes related to body fat distribution.

## Results

### Suggestive archaic ancestry in Greenlandic Inuit SNP chip data

We first computed *f_D_* (Martin, et al. 2014) to assess putative local archaic human ancestry in candidate genes in the highest 99.5% quantile of the PBS genome-wide distribution in GI (Fumagalli, et al. 2015). This statistic is intended to detect imbalances in the sharing of alleles from an outgroup panel between two sister population panels or genomes. It is robust to differences in local diversity along the genome, which tends to confound other related statistics, such as Patterson’s D^14,15^, which also use patterns of allelic imbalances (i.e. “ABBA” and “BABA” sites, see Methods). Patterson’s *D* is also meant to identify excess archaic ancestry but is better suited for genome-wide analyses (Green, et al. 2010; Durand, et al. 2011).

For the test population we used the SNP chip data from the selection scan in GI (Fumagalli, et al. 2015), obtained from 191 Greenlandic Inuits with low European admixture (<5%). When computing *f_D_*, we used a Yoruba genome sequenced to high-coverage (HGDP00927) (Prüfer, et al. 2014) as the non-introgressed genome. For the archaic genome, we used either a high-coverage Denisovan genome (Meyer, et al. 2012) or a high-coverage Neanderthal genome (Prüfer, et al. 2014). All sites were polarized with respect to the inferred human-chimpanzee ancestral state (Paten, et al. 2008).

The only top PBS locus showing some evidence of archaic ancestry is the *WARS2/TBX15* region, which has a high degree of allele sharing with the Denisovan genome (Figures S1, S2, Table S1). For example, in *WARS2*, we find 18.46 sites supporting a local tree in which Denisova is a sister group to GI, to the exclusion of Yoruba (ABBA), while we only find 2.5 sites supporting a Yoruba-Denisova clade, to the exclusion of GI (BABA) (the numbers are not integers because we are using the panel version of *ABBA* and *BABA*, not the single-genome version (Durand, et al. 2011)). However, because we used SNP chip data, we found that most genes had few informative (ABBA or BABA) sites. For genes for which there are 2 or more informative sites (i.e. ABBA+BABA >= 2), *TBX15* is in the 88% quantile of the genome-wide distribution of *f_D_*, and *WARS2* is in the 87% quantile. For genes for which there are more than 10 informative sites (i.e. ABBA+BABA >= 10), the quantile rankings for these genes are almost the same: *TBX15* is in the 88% quantile, and *WARS2* is in the 86% quantile. However, due to the small number of SNPs available for testing in GI, we could not assess with confidence whether this region was truly introgressed from archaic humans. We therefore sought to identify the selected haplotype in other samples for which full sequencing data are available.

The alleles with high PBS values and high frequency in GI are almost absent in Africa, while present across Eurasia. In Figure 1.A, we show the geographic distribution of allele frequencies for one of these SNPs (rs2298080) as an example, using data from phase 3 of the 1000 Genomes Project (Auton, et al. 2015) and the Geography of Genetic Variants Browser (Marcus and Novembre 2016). The high-frequency alleles in GI tend to match the Denisovan and Altai Neanderthal alleles in this region. For example, rs2298080 has an A allele at a frequency of 45.45% in Han Chinese from Beijing (CHB) and at 99.74% frequency in GI. This allele is absent or almost absent (<1% frequency) in all African populations, and the Denisovan and Altai Neanderthal genomes are both homozygous for the A allele. We observe a similar pattern when looking at the Simons Genome Diversity Project (SGDP) (Figure 1.B), which contains high-coverage genomes from a wide variety of populations across the world, including San, native Papuans and Australians, and various Native American populations. Here, we also observe that the archaic allele of rs2298080 is almost absent (1.11% frequency) in Africans, but has a much higher frequency outside the African continent, especially in East Asia (46.36%), Central Asia and Siberia (64.81% frequency), and the Americas (89.13% frequency), though it is not at very high frequencies in Oceania (20.37% frequency). We therefore analyze sequencing data from Eurasians to determine if the selected alleles were truly introgressed from an archaic human population.

**Figure 1.**
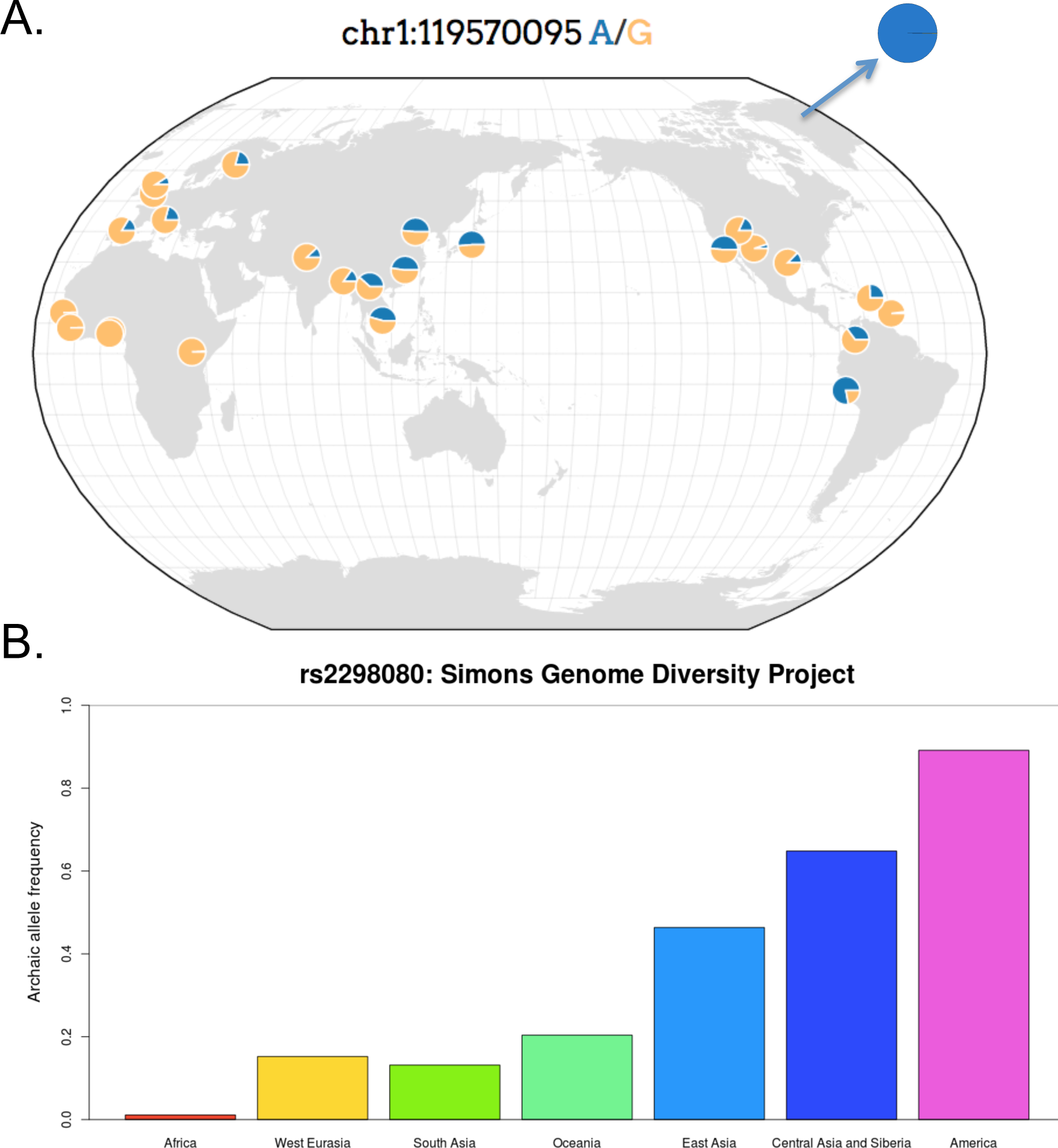
A. Geographic distribution of rs2298080 in different 1000 Genomes populations. The color blue corresponds to the archaic allele in this SNP. For comparison, the allele frequency in Greenlandic Inuit is also shown. This figure was made using the Geography of Genetic Variants browser v.0.1, by J. Novembre and J.H. Marcus: http://popgen.uchicago.edu/ggv/. B. Archaic allele frequencies of rs2298080 in the continental panels of the Simons Genome Diversity Project.

### Excess archaic ancestry in Eurasians

A particularly useful way to detect adaptive introgression is to identify regions with a high proportion of uniquely shared sites between the archaic source population and the population subject to introgression (Racimo, et al. 2016). This was one of the lines of evidence in favor of Denisovan adaptive introgression in Tibetans at the *EPAS1* locus (Huerta-Sánchez, et al. 2014). We therefore partitioned the genome into non-overlapping 40 kb windows and computed, in each window, the number of SNPs where the Denisovan allele is at a frequency higher than 20% in Eurasians but less than 1% in Africans, using the populations panels from phase 3 of the 1000 Genomes Project (Auton, et al. 2015). The windows containing *TBX15* and *WARS2* have four and three such sites, respectively, which is higher than 99.99% of all windows in the genome (Figure 2.A). We used a length of 40 kb because the mean length of introgressed haplotypes found in an earlier study (Prüfer, et al. 2014) was 44,078 bp (Supplementary Information 13 in that study).

**Figure 2.**
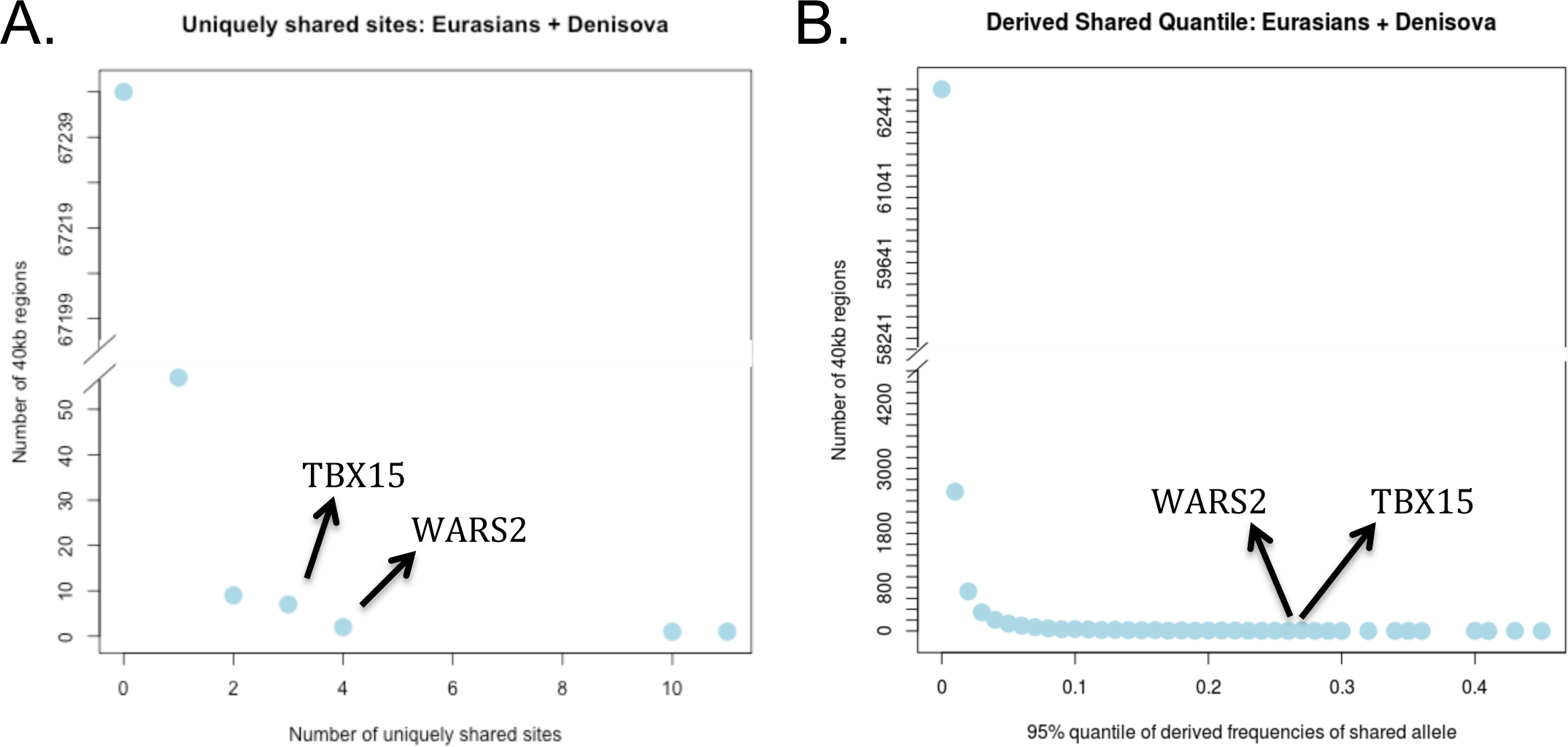
A) Genome-wide histogram of the number of uniquely shared sites where the Denisovan allele is at less than 1% frequency in Africans (AFR, excluding admixed African-Americans) and at more than 20% frequency in Eurasians (EUR + SAS + EAS). The counts were computed in non-overlapping 40 kb regions of the genome. The y-axis is truncated, as the vast majority of regions have 0 uniquely shared sites. *TBX15* and *WARS2* are among the few regions that have 3 and 4 uniquely shared sites, respectively. B) In each of the same 40 kb windows, we also computed the 95% quantile of Eurasian derived allele frequencies of all SNPs that are homozygous derived in Denisova and less than 1% derived in Africans. The figure shows a histogram of this score for all windows. The 95% quantile scores for *TBX15* and *WARS2* are 0.27 and 0.26, respectively.

In each of the same 40 kb windows, we also computed the 95% quantile of Eurasian derived allele frequencies of all SNPs that are homozygous derived in Denisova and less than 1% derived in Africans (using EPO alignments (Paten, et al. 2008) to define the ancestral and derived alleles with respect to the human-chimpanzee ancestor). This quantile is assigned as the score for each window. This second statistic is designed to detect archaic alleles that are uniquely shared with Eurasians and have risen to extremely high frequencies. Here, *WARS2* and *TBX15* are also strong outliers, with a quantile frequency score above 99.95% of all windows (Figure 2.B). The chance that the region would randomly show such an extreme pattern of both excess number, and high allele frequency, of derived alleles shared with Denisovans and rare or absent in Africa, is exceedingly small under models of neutrality, positive selection alone or introgression alone (Racimo, et al. 2016) and strongly suggests that selection has been acting on alleles introgressed from archaic humans.

### Identifying the introgression tract

We used a Hidden Markov Model (HMM) method (Seguin-Orlando, et al. 2014) for identifying introgression tracts (“HMM-tracts”), using Yoruba as the population without any introgression (Supplementary Information 1). Using either Neanderthal or Denisova as the source population, we inferred a clear introgression tract in the middle of the PBS peak, which is especially frequent among East Asians and Americans (Figure S3). The inferred tract is more than twice as wide when using Denisova (chr1:119,549,418-119,577,410) as the source population (27,992 bp) than when using Neanderthal (13,729 bp) (Figure S4), which suggests the source population may have been more closely related to the Denisovan genome in this particular region. The frequency of the Denisovan tract is 48.41% in East Asians, 19.78% in Europeans and 12.37% in South Asians. We find 5 individuals that contain particularly long (129,374 bp to 216,155 bp) versions of the tract in East Asians: one Southern Han Chinese (HG00406), two Kinh (HG01841, HG02076) and three Japanese (NA18962, NA19005, NA19054).

We also queried the region using the output from the conditional random field framework for detecting archaic tracts developed by Sankararaman et al. (2016), applied to the Simons Genome Diversity Panel. Using this method (“CRF-tracts”), we find that the tract is generally longer (~85,000 bp, though varying in length depending on the population) than the one inferred by the HMM, for example extending in several East Asians from position 119,541,452 to 119,627,434 (Figure 3). We also find that there are individuals from three populations in East and Central Asia (the Naxi, the Yakut and the Even) in which the tract is surprisingly much larger than in any other population (ranging from 181,288 bp to 628,816 bp). Although the Yakut are Siberian populations that are closely related to the Inuit, the version of the introgressed haplotype that is at high frequency in the Yakut does not appear to match better with the Inuit haplotype than other versions of the introgressed haplotype present in other populations (Supplementary Information 2).

**Figure 3.**
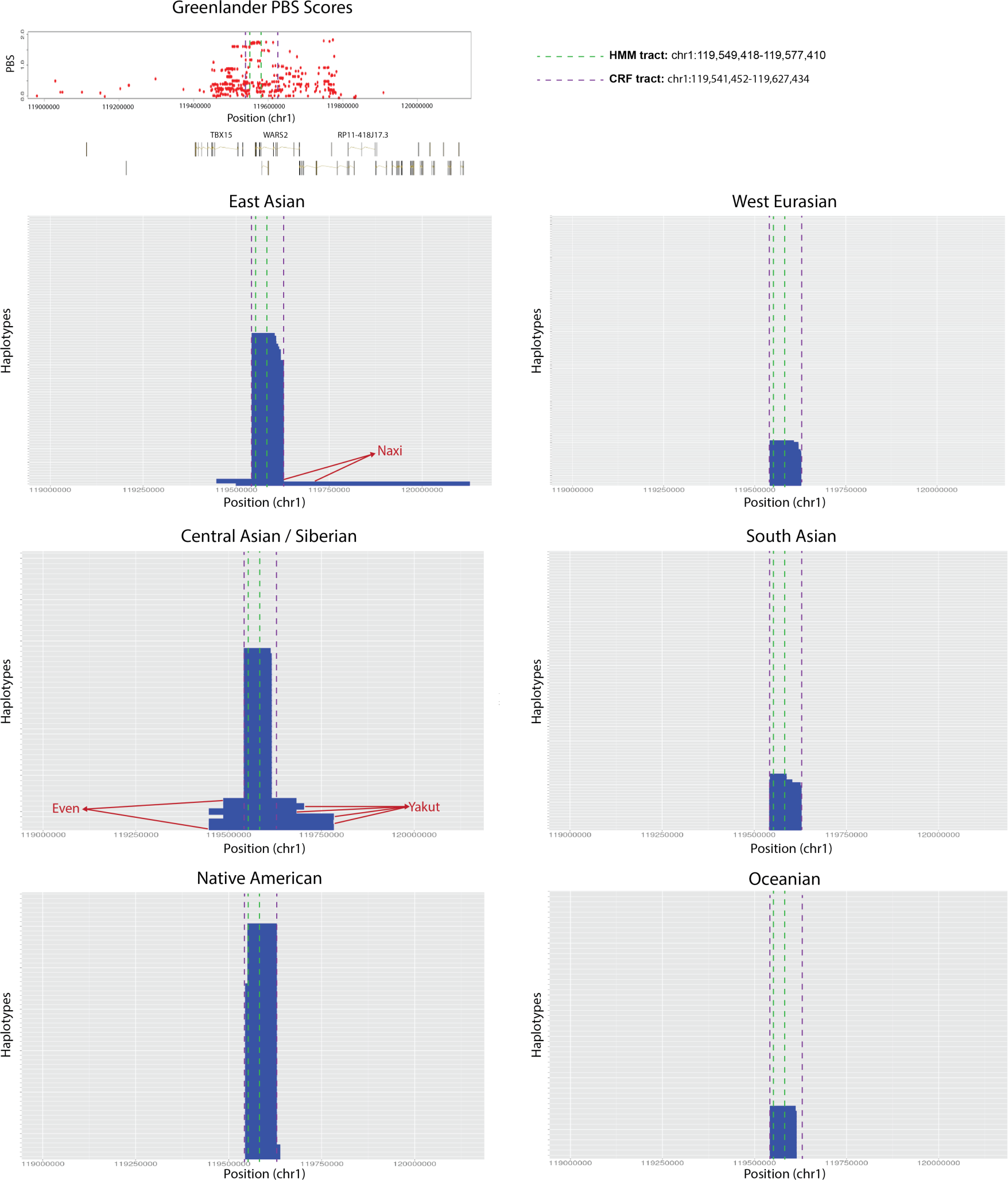
Introgression tracts inferred from the CRF method (Sankararaman et al. 2016) in the different continental panels of the SGDP data, using Denisova as the source population. The tract (denoted by the purple dashed line) coincides with the peak of PBS scores obtained in the GI scan for positive selection (red dots). For comparison, we also show the boundaries of the tract as inferred by the HMM method (green dashed line). The Even, Yakut and Naxi contain considerably longer versions of the tract. The frequency of the introgressed haplotype is highest among Native Americans, but is also at intermediate frequencies in Central and East Asians, and Siberians.

We further examined the SNPs that are selected in GI and that contain archaic alleles that are uniquely present in the introgressed haplotype background, as inferred by the HMM. We queried their allelic state in different human population panels (Figure 4). We find a sharp distinction between the two most prevalent haplotypes across Eurasia, with one of them being almost identical to the Denisovan genome and the other being highly differentiated from it. As expected, the frequencies of these two haplotypes agree with the frequencies of the inferred introgressed tracts and with the allelic frequencies of the top PBS SNPs. This pattern echoes the pattern observed for another well-known case of adaptive introgression in Tibetans (Huerta-Sánchez, et al. 2014), although in the present case the archaic haplotype is more widely distributed across Eurasia.

**Figure 4.**
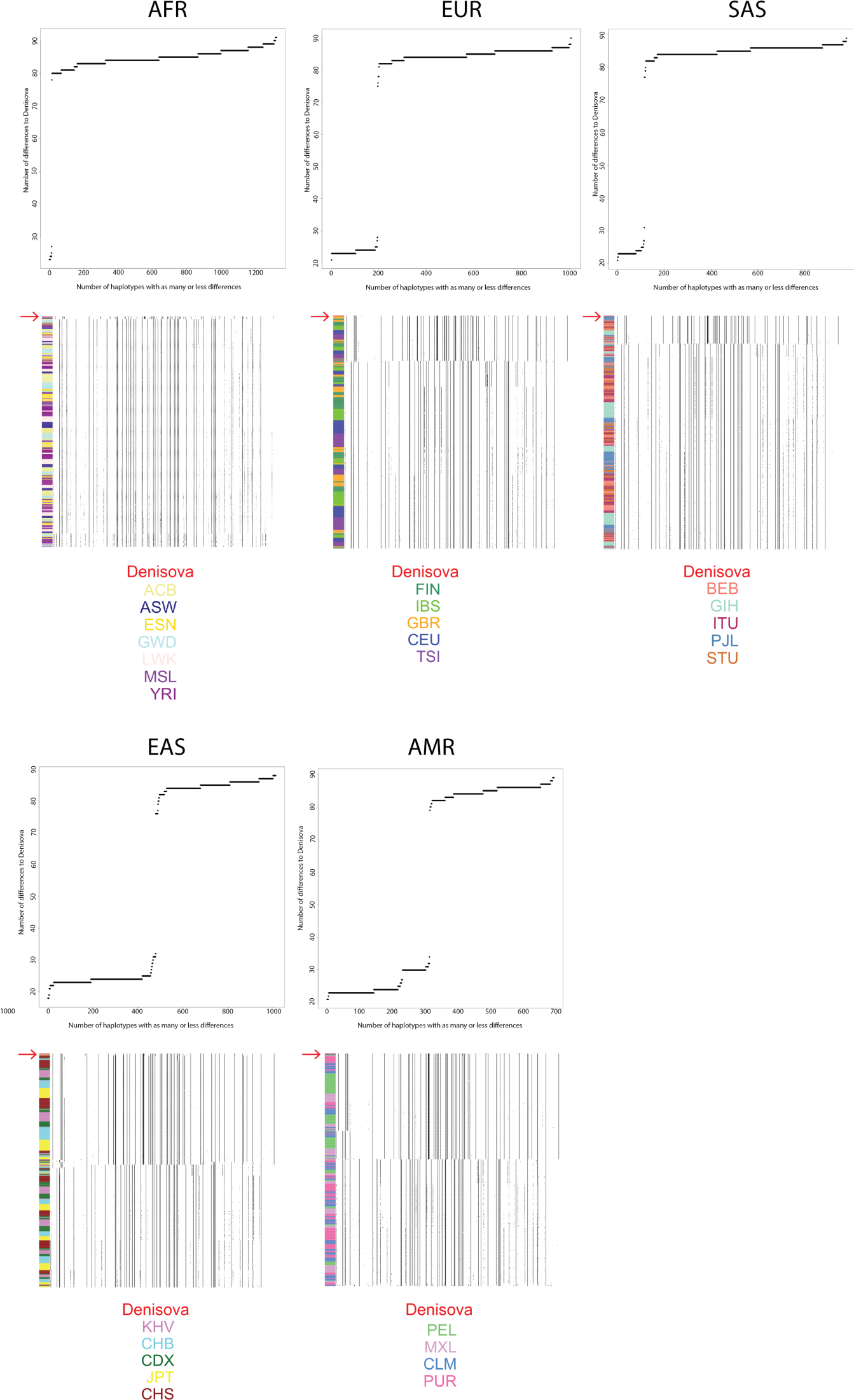
We counted the differences between the Denisovan genome and all haplotypes in each 1000G continental panel (AFR, EUR, SAS, EAS and AMR). For each panel, we plot the cumulative number of haplotypes that have as many or less differences to the Denisovan genome than specified in the x-axis. Below each of these plots, we also plot the haplotype structure for each 1000G panel in the introgressed tract, ordering haplotypes by decreasing similarity to the Denisovan genome (red arrow, at the top of each panel). The color codes at the bottom refer to 1000G sub-populations to which the different haplotypes belong, as indicated by the right color column in each panel.

The average recombination rate per bp in the region is low (0.15 * 10^-8, Figure S5), so we were interested in testing if the inferred tract could possibly be an ancestral polymorphism that survived in particular populations but not others, rather than an introgressed haplotype. Using the observed recombination rate and the conservative, shorter tract length inferred by the HMM (27,992 bp), we find via simulations that this probability is very small (P = 0.01, Supplementary Information 3). Additionally, we note that this probability would be even lower using the tract length inferred by the CRF method, or the much longer tracts found in the Yakut, Even and Naxi.

### The selected alleles are the putative introgressed alleles

The archaic haplotype frequencies agree well with the frequencies of the selected alleles in different human populations. We therefore aimed to verify that the selected alleles were the same alleles that were uniquely shared with archaic humans. We focused on the SNPs in the *TBX15/WARS2* region that are located in the 99.95% quantile of genome-wide PBS scores in the GI SNP chip data (Fumagalli, et al. 2015). Noticeably, 28 out of the 33 top SNPs in the *TBX15/WARS2* region lie in the region where the introgressed haplotype is located, as inferred by both the HMM and the CRF frameworks. Out of the remaining 5 SNPs, two of them (rs10923738, rs12567111) lie within the inferred CRF-tract, but not in the HMM-tract. For each of SNPs that overlap the track as inferred by both methods, we checked whether the selected allele in GI was the same as: a) the alleles present in the high-coverage Altai Neanderthal genome(Prüfer, et al. 2014), b) the alleles present in the recently-sequenced high-coverage Vindija Neanderthal genome (https://bioinf.eva.mpg.de/jbrowse), c) the alleles in the high-coverage Denisova genome (Meyer, et al. 2012), d) the alleles present in a present-day human genome from the 1000 Genomes Project(Auton, et al. 2015) (HG00436) that is homozygous for the introgressed tract, e) the alleles in a present-day human genome (HG00407) that does not contain the introgressed tract, and f) the alleles in 3 modern human genomes obtained from ancient DNA: Ust-Ishim (Fu, et al. 2014) (dated at ~45,000 kya), Stuttgart (dated at ~7,000 kya) and Loschbour (Lazaridis, et al. 2014) (dated at ~8,000 kya).

In all of the SNPs showing the highest evidence of selection, the present-day human genomes that are homozygous for the introgression tract are also homozygous for the favored alleles. Additionally, in all of these SNPs, the present-day human genomes lacking the introgression tract are homozygous for the allele that was not favored by selection. Furthermore, in 79% of the SNPs, the selected allele is present in homozygous form in Denisova, while this is only true for 64% of the SNPs in Neanderthals (Table 1), regardless of whether we look at the high-coverage Altai Neanderthal or at the recently sequenced high-coverage Vindija Neanderthal. This indicates that the selected alleles are also the introgressed alleles, and that a population closer to the sequenced Denisovan at this locus is the most likely source of archaic introgression. In the 6 SNPs where the introgressed tract carries a different allele than Denisova, the allele in the introgressed tract is derived, suggesting these differences are due to mutations that occurred more recently than the time the introgressing lineage coalesced with the sequenced Denisovan’s lineages, and possibly more recently than the introgression event. Finally, we find that in 100% of the SNPs the Ust’-Ishim, Stuttgart and Loschbour genomes are homozygous for the non-introgressed alleles, so they did not carry the introgressed haplotype.

**Table 1.** The 28 SNPs in the 99.95% highest PBS quantile in Greenlandic Inut that lie in the introgressed tract as determined by the HMM. CHR = chromosome. POS = position (hg19). SNP ID = dbSNP rs ID number. REF = reference allele. ANC = human-chimpanzee ancestor allele (based on EPO alignments). NONSEL = non-selected allele. SEL = selected allele. GR FREQ = frequency of the selected allele in Greenlandic Inuit. CHB FREQ = frequency of the selected allele in CHB (Chinese individuals from Beijing). CEU FREQ = frequency of the selected allele in CEU (Individuals of Central European descent living in Utah). ALTAI = selected allele counts in the high-coverage Altai Neanderthal genome. VIN = selected allele counts in the high-coverage Vindija Neanderthal genome (https://bioinf.eva.mpg.de/jbrowse/). DEN = selected allele counts in the Denisova genome. HG00436 = selected allele counts in a present-day human genome that is homozygous for the introgressed tract. HG00407 = selected allele counts in a present-day human genome that is homozygous for the absence of the tract. UST’-ISHIM = selected allele counts in the Ust’-Ishim genome. STUTTGART = selected allele counts in the Stuttgart genome. LOSCHBOUR = selected allele counts in the Loschbour genome.

| CHR | POS | SNP ID | REF | ANC | NONSEL | SEL | GR FREQ | CHB FREQ | CEU FREQ | ALTAI | VIN | DEN | HG00436 | HG00407 | UST'-ISHIM | STUTTGART | LOSCHBOUR |
| --- | --- | --- | --- | --- | --- | --- | --- | --- | --- | --- | --- | --- | --- | --- | --- | --- | --- |
| 1 | 119570095 | rs2298080 | G | A | G | A | 0.997382 | 0.4545 | 0.1833 | 2 | 2 | 2 | 2 | 0 | 0 | 0 | 0 |
| 1 | 119571463 | rs10923735 | C | T | C | T | 0.997382 | 0.4545 | 0.1833 | 2 | 2 | 2 | 2 | 0 | 0 | 0 | 0 |
| 1 | 119574289 | rs12030972 | G | G | G | A | 0.997382 | 0.4545 | 0.1833 | 0 | 0 | 0 | 2 | 0 | 0 | 0 | 0 |
| 1 | 119574537 | rs12026409 | G | G | G | A | 0.997382 | 0.4545 | 0.1833 | 0 | 0 | 0 | 2 | 0 | 0 | 0 | 0 |
| 1 | 119557151 | rs10923726 | A | A | A | G | 0.997382 | 0.4659 | 0.1833 | 0 | 0 | 2 | 2 | 0 | 0 | 0 | 0 |
| 1 | 119557772 | rs4659140 | T | T | T | C | 0.997382 | 0.4659 | 0.1833 | 0 | 0 | 0 | 2 | 0 | 0 | 0 | 0 |
| 1 | 119558112 | rs12027501 | C | C | C | G | 0.997382 | 0.4659 | 0.1833 | 0 | 0 | 0 | 2 | 0 | 0 | 0 | 0 |
| 1 | 119558126 | rs12027524 | C | T | C | T | 0.997382 | 0.4659 | 0.1833 | 2 | 2 | 2 | 2 | 0 | 0 | 0 | 0 |
| 1 | 119559155 | rs12410034 | A | T | A | T | 0.997382 | 0.4659 | 0.1833 | 2 | 2 | 2 | 2 | 0 | 0 | 0 | 0 |
| 1 | 119559260 | rs4659141 | T | T | T | G | 0.997382 | 0.4659 | 0.1833 | 0 | 0 | 0 | 2 | 0 | 0 | 0 | 0 |
| 1 | 119559297 | rs4658995 | C | C | C | T | 0.997382 | 0.4659 | 0.1833 | 2 | 2 | 2 | 2 | 0 | 0 | 0 | 0 |
| 1 | 119559607 | rs1325932 | C | T | C | T | 0.997382 | 0.4659 | 0.1833 | 2 | 2 | 2 | 2 | 0 | 0 | 0 | 0 |
| 1 | 119559793 | rs1325931 | A | C | A | C | 0.997382 | 0.4659 | 0.1833 | 2 | 2 | 2 | 2 | 0 | 0 | 0 | 0 |
| 1 | 119559796 | rs1325930 | A | A | A | C | 0.997382 | 0.4659 | 0.1833 | 2 | 2 | 2 | 2 | 0 | 0 | 0 | 0 |
| 1 | 119560089 | rs10923730 | C | C | C | G | 0.997382 | 0.4659 | 0.1833 | 2 | 2 | 2 | 2 | 0 | 0 | 0 | 0 |
| 1 | 119560512 | rs963171 | A | T | A | T | 0.997382 | 0.4659 | 0.1833 | 2 | 2 | 2 | 2 | 0 | 0 | 0 | 0 |
| 1 | 119561611 | rs3951792 | G | G | G | A | 0.997382 | 0.4659 | 0.1833 | 0 | 0 | 2 | 2 | 0 | 0 | 0 | 0 |
| 1 | 119562014 | rs12024063 | G | G | G | A | 0.997382 | 0.4659 | 0.1833 | 0 | 0 | 0 | 2 | 0 | 0 | 0 | 0 |
| 1 | 119562180 | rs12031562 | C | C | C | G | 0.997382 | 0.4659 | 0.1833 | 0 | 0 | 2 | 2 | 0 | 0 | 0 | 0 |
| 1 | 119562274 | rs12031597 | C | C | C | T | 0.997382 | 0.4659 | 0.1833 | 2 | 2 | 2 | 2 | 0 | 0 | 0 | 0 |
| 1 | 119562382 | rs12021830 | T | C | T | C | 0.997382 | 0.4659 | 0.1833 | 2 | 2 | 2 | 2 | 0 | 0 | 0 | 0 |
| 1 | 119562460 | rs10923733 | A | A | A | G | 0.997382 | 0.4659 | 0.1833 | 2 | 2 | 2 | 2 | 0 | 0 | 0 | 0 |
| 1 | 119562557 | rs10494218 | A | T | A | T | 0.997382 | 0.4659 | 0.1833 | 2 | 2 | 2 | 2 | 0 | 0 | 0 | 0 |
| 1 | 119562716 | rs10494219 | T | C | T | C | 0.997382 | 0.4659 | 0.1833 | 2 | 2 | 2 | 2 | 0 | 0 | 0 | 0 |
| 1 | 119562920 | rs12402563 | G | G | G | C | 0.997382 | 0.4659 | 0.1833 | 0 | 0 | 2 | 2 | 0 | 0 | 0 | 0 |
| 1 | 119562932 | rs12405154 | C | C | C | T | 0.997382 | 0.4659 | 0.1833 | 2 | 2 | 2 | 2 | 0 | 0 | 0 | 0 |
| 1 | 119563172 | rs12025128 | G | A | G | C | 0.997382 | 0.4659 | 0.1833 | 2 | 2 | 2 | 2 | 0 | 0 | 0 | 0 |
| 1 | 119563638 | rs113389819 | G | C | G | C | 0.997382 | 0.4659 | 0.1833 | 2 | 2 | 2 | 2 | 0 | 0 | 0 | 0 |

Additionally, by analyzing low-coverage ancient DNA data from various Eurasian populations (Gamba, et al. 2014; Raghavan, et al. 2014; Allentoft, et al. 2015), we find that the selected alleles are prevalent in Eurasian steppe populations but almost absent in western Neolithic and Mesolithic European populations (Supplementary Information 4), suggesting the introgressed haplotype may have been introduced into Europe from eastern Eurasia, via the Late Neolithic steppe migrations (Allentoft, et al. 2015; Haak, et al. 2015).

### The region also shows selection signatures in Native Americans

To examine whether selection in GI on the introgressed haplotype was shared with Native Americans, we performed a scan for positive selection in the latter population by measuring the genetic differentiation against a population of African descent. To correct for recent admixture in Latin America, we selected the Latin American individuals from the 1000 Genomes project (Auton, et al. 2015) showing the highest proportion of Native American ancestry in a 100 Mbp region around the introgressed haplotype, using *ngsAdmix* (Skotte, et al. 2013). These came only from individuals from Peru (PEL) and from individuals from Los Angeles with Mexican ancestry (MXL), as these have higher proportions of Native American ancestry than other Latin American panels (Figure S6).

We observed a local increase of *F_ST_* in the proximity of the introgressed haplotype when comparing individuals from the PEL and MXL panels against populations of African (YRI) descent (Figure 5). We did not observe high *F_ST_* values when testing panels of East Asian, South Asian or European descent as the target population for selection (Figure S7).

**Figure 5.**
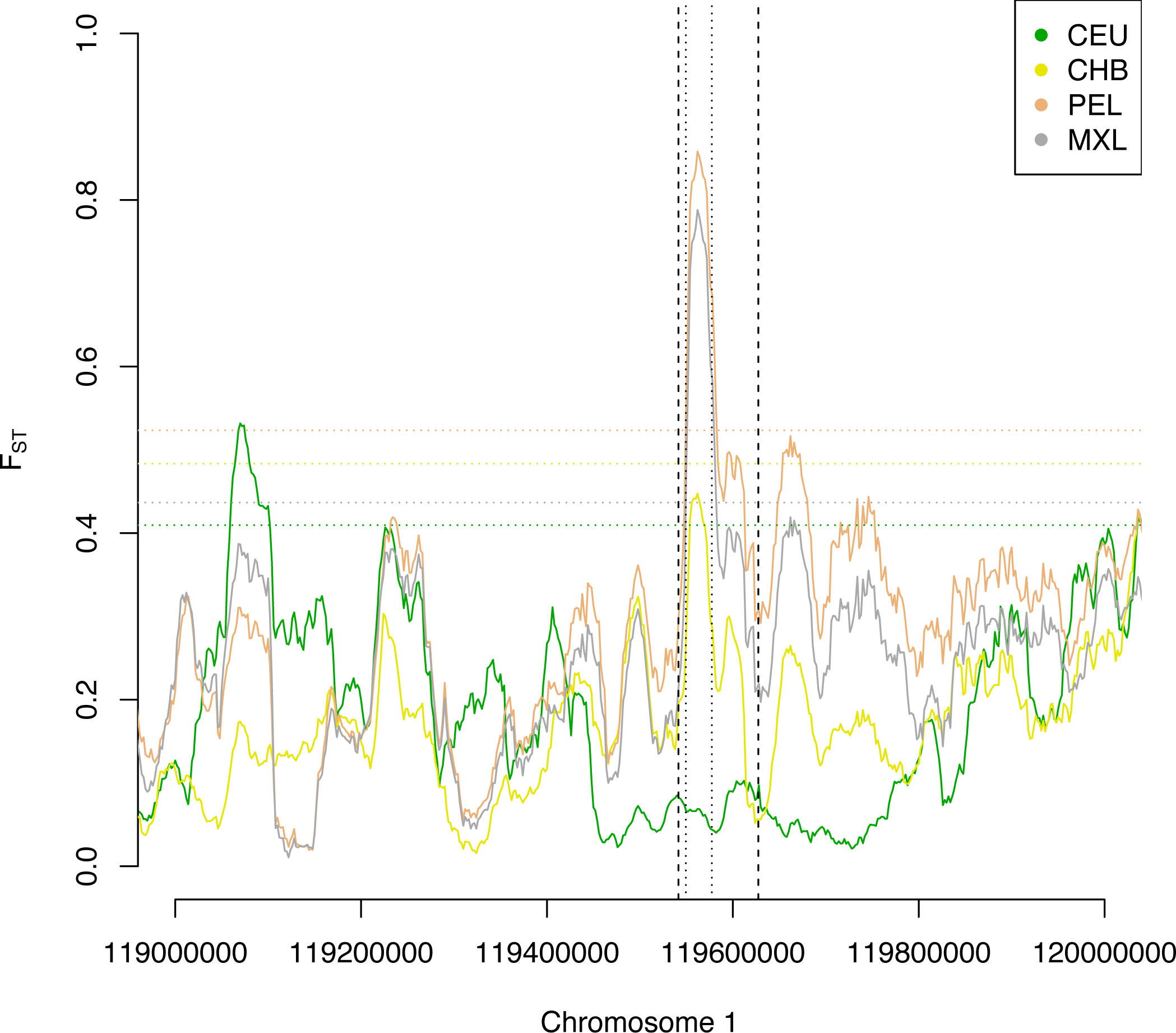
We computed *F_ST_* in different non-African populations (CEU, CHB, PEL, MXL) against African Yoruba (YRI) in the *TBX15/WARS2* region. We find a significant (P < 0.01) local increase in *F_ST_* in American populations (PEL, MXL) exactly where the introgressed haplotype is inferred to be (CRF-tract: vertical dashed lines, HMM-tract: vertical dotted lines). In contrast, we found no significant increase in *F_ST_* when comparing Europeans (CEU) and East Asians (CHB) against YRI (although there is an observable but non-significant peak in CHB). 99th percentile values of the *F_ST_* empirical distribution for all comparisons are represented as horizontal dotted lines and color-coded according to the corresponding population.

To assess whether the observed values of *F_ST_* around the introgressed haplotype may be explained by pure genetic drift, we calculated *F_ST_* for all biallelic SNPs at the whole-genome level and compared the empirical distribution with a marker SNP (rs2298080) for the selected haplotype. We found that *F_ST_* values between PEL/MXL against YRI or CEU for the introgressed haplotype are strong outliers in the genome-wide distribution (Table S2). *F_ST_* between PEL and CHB is also an outlier (p=0.021) while *F_ST_* between CHB and CEU is marginally significant (p=0.043).

We aimed to elucidate whether the high *F_ST_* values observed between East Asians and Native Americans could be due solely to selection in the ancestral population of Native Americans and GI, or if they could be better explained by continuing selection in Native Americans after their split from GI. We simulated several demographic scenarios of the history of CHB, PEL and GI, under various PEL-GI split times and PEL population sizes (see Methods). We then tested whether the top *F_ST_* for our 20kbp sliding window scan was an outlier under scenarios with positive selection starting in the ancestral population of PEL and GI, and either continuing in both PEL and GI, or only persisting in GI. We found that demographic scenarios involving low effective population sizes in PEL and/or very strong selection in the ancestral population could result in the observed *F_ST_* value being in the 95% percentile of the distribution, without invoking continuing selection in Native Americans after their split from GI (Figure S8). The population sizes required for this to occur (1,000-3,000) overlap with previously estimated population size estimates for Native Americans (1,650-2,600)(Raghavan, et al. 2015), so it is possible that selection need not have continued to operate in Native Americans after their split from GI.

Using rs2298080 as a proxy for the selected haplotype, we infer that the haplotype has a frequency of 98%, and 86% in the PEL and MXL population panels, after filtering for the individuals with the most Native American ancestry, respectively. The apparent lack of fixation of the introgressed haplotype in these populations might be explained by the residual non-Native American ancestry (ranging from 11% to 29%) for the analyzed American individuals. Clearly, selection on this allele is not unique to GI but has affected a large proportion of New World groups, likely in pre-Columbian times. This also explains the higher frequency of the introgressed tract in the SGDP Native American panel than in the 1000 Genomes Project panel (Figures 1.B, 3), as the individuals from the former project were specifically sampled from peoples with well-documented Native American ancestry.

### The introgression tract is more closely related to the Denisovan genome than the Neanderthal genome

The divergence between the introgressed haplotype (as defined by the HMM) and the Denisova genome (0.0008) is lower than the divergence between the haplotype and the Altai Neanderthal genome (0.0016). To further examine if the haplotype could be of Neanderthal origin, we computed the divergence between the Altai Neanderthal genome and a randomly chosen individual that was homozygous for the introgressed tract (HG00436). We compared this divergence to the distribution of divergences between the Altai Neanderthal and the Mezmaiskaya Neanderthal (Prüfer, et al. 2014), computed across windows of the genome of equal size (Figure S9). We scaled all divergences by the divergence between the human reference and the inferred human-chimpanzee ancestor, to account for local variation in mutation rates. If the introgressed haplotype came from Neanderthals, then we would expect the scaled divergence between it and the Altai Neanderthal to be within the distribution of Neanderthal-Neanderthal divergence. To avoid errors due to the lower coverage of the Mezmaiskaya Neanderthal, we only computed divergence on the Altai side of the tree. The observed Altai-haplotype scaled divergence falls in the 96.9% quantile of the distribution, suggesting it is likely not a typical Neanderthal haplotype. The divergence between the introgressed haplotype and the Denisovan genome falls in the 43.12% quantile of the Neanderthal-Denisovan divergence distribution, indicating that even though the haplotype is more closely related to the sequenced Densovan than to the Altai Neanderthals, it is just as diverged to the Denisovan as a typical Neanderthal region is to a typical Denisovan region.

We also obtained the distribution of scaled divergences between the Denisovan genome and a high-coverage Yoruba genome (HGDP00936)(Prüfer, et al. 2014). We compared this distribution to the divergence between the introgressed haplotype (using an individual homozygous for the haplotype), and the Denisovan, and observe that this divergence falls towards the left end of the distribution, in the 16.44% quantile (Figure S9). Furthermore, the introgressed haplotype is highly diverged from Yoruban haplotypes, with the divergence falling in the 98.5% quantile of the Denisova-Yoruba divergence distribution, and in the 98.67% quantile of the Neanderthal-Yoruba divergence distribution (Figure S9). All in all, this suggests that both the introgressed haplotypes and the archaic haplotypes – though closely related to each other – are both highly diverged from the non-introgressed present-day human haplotypes.

Additionally, we simulated two archaic populations that split at different times (100,000, 300,000 or 450,000 years), under a range of effective population sizes (1,000, 2,500, 5,000), based on estimates from an earlier study (Prüfer, et al. 2014). We compared these simulations to the divergence between the Neanderthal genome and the introgressed haplotype, as well as to the divergence between the Denisovan genome and the introgressed haplotype and the divergence between the introgressed haplotype and the Yoruba genome (Figure S10). In all cases, the divergence that falls closest to the distribution is the divergence to the Denisovan genome, suggesting this is the closest source population for which we have sequence data.

To understand the relationship between the introgressed haplotype and the archaic and present-day human genomes, we plotted a network of the haplotypes in the region as defined by the HMM method (Figure 6). This network shows the most parsimonious distances among the 20 most common present-day human haplotypes and the 4 archaic haplotypes from Altai Neanderthal and Denisova in the region. We observe two distinct clusters of present-day human haplotypes: one cluster that is distantly related to both archaic genomes, and another cluster that is closely related to Denisova and contains mostly East Asian and Native American individuals, some European and South Asian individuals and almost no Africans. The Altai Neanderthal haplotypes fall at an intermediate position between the Denisovan haplotypes and the first cluster, but share more similarities with the Denisovan haplotypes. The present-day human cluster that is closest to Denisova has a smaller Hamming distance to Denisova than does the Neanderthal haplotype (Figure S11), again suggesting the haplotype was introgressed from individuals more closely related to the Denisovan genome in this region than to any of the sequenced Neanderthal genomes.

**Figure 6.**
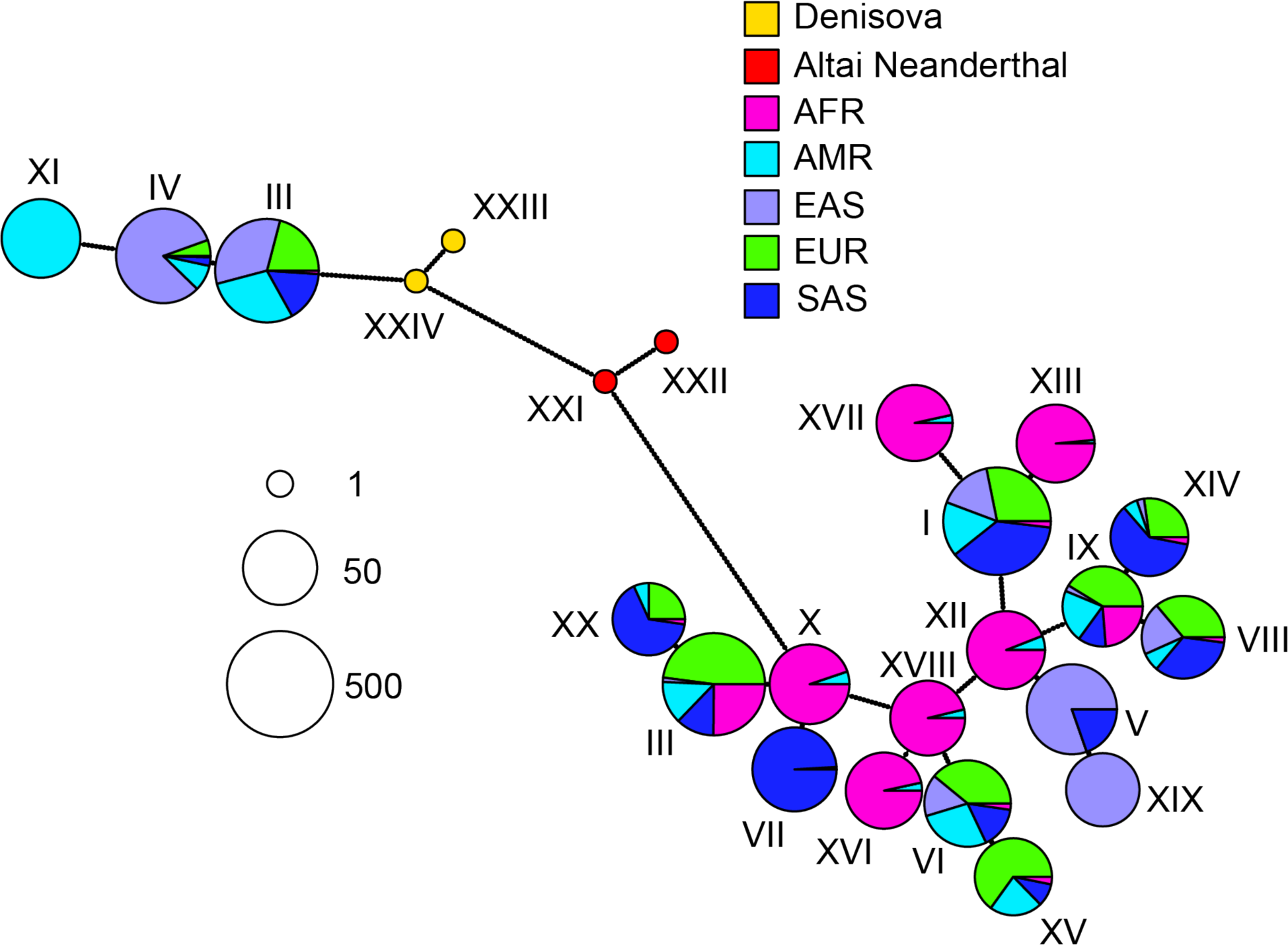
Network of archaic haplotypes and 20 most common present-day human haplotypes from the 1000 Genomes Project in the *TBX15/WARS2* introgressed region as inferred by the HMM. Each pie chart is a haplotype, and the dots along each line represent the number of differences between each haplotype. The size of each pie chart is proportional to the log base 2 of the number of individuals in which that haplotype appears, and the colors refer to the proportion of those individuals that come from different continental populations. AFR: Africans. AMR: Americans. EAS: East Asians. EUR: Europeans. SAS: South Asians.

We compared the distances between the haplotypes in *TBX15/WARS2* to those observed in *EPAS1* (Figure S12), a previously reported case of adaptive introgression in Tibetans from an archaic population closely related to the Denisovan genome(Huerta-Sánchez, et al. 2014; Huerta-Sanchez and Casey 2015). We focused on the distances among Neanderthal, Denisova and the haplotype that is most closely related to the archaic genomes, for each of the distinct present-day clusters. We observe that, although the distances among Denisova, Neanderthal and the putatively introgressed haplotypes are similar, the distance between any of these and the non-introgressed haplotype are approximately double of what is observed in the case of *EPAS1* (Figure S12). The mutation rate in the region does not seem particularly high, based on the patterns of divergence between the human reference and the human-chimpanzee ancestor (Figure S13), so a locally elevated mutation rate is unlikely to explain this signal. This also suggests that the archaic and introgressed haplotypes in this region belong to a deeply divergent archaic lineage.

### Regulatory differences in archaic and modern humans

We previously identified four regions in *TBX15* in which the Denisovan DNA methylation patterns significantly differ from those of present-day humans (Gokhman, et al. 2014). This high concentration of differentially methylated regions (DMRs) makes *TBX15* one of the most DMR-rich genes in the Denisovan genome. These DMRs are found around the transcription start site (TSS) of *TBX15*, bear many active chromatin marks in present-day humans (DNase I hyper-sensitivity, binding by p300 and H2A.Z, and the histone modifications H3K27ac and H3K4me1), and were shown to be associated with the activity levels of *TBX15* (Kron, et al. 2012; Chandra, et al. 2014). This suggests that the activity level of *TBX15* in the Denisovan genome was different than in present-day human genomes.

The extensive differences in DNA methylation between the Denisovan and present-day human genomes, and the fact that the introgressed tract overlaps regulatory regions of *TBX15* (Chandra, et al. 2014) suggest that present-day individuals who carry the introgressed haplotype might display differential methylation as well. Despite the fact that the four Denisovan DMRs do not overlap the introgressed region (Figure 7A), they might reflect *TBX15* activity levels determined by sequence changes at the introgressed region, as it is not uncommon that sequence changes far from a gene alter its activity (Mansour, et al. 2014; Diederichs, et al. 2016). Thus, we turned to investigate the link between the introgressed allele and DNA methylation around *TBX15* and *WARS2*.

**Figure 7.**
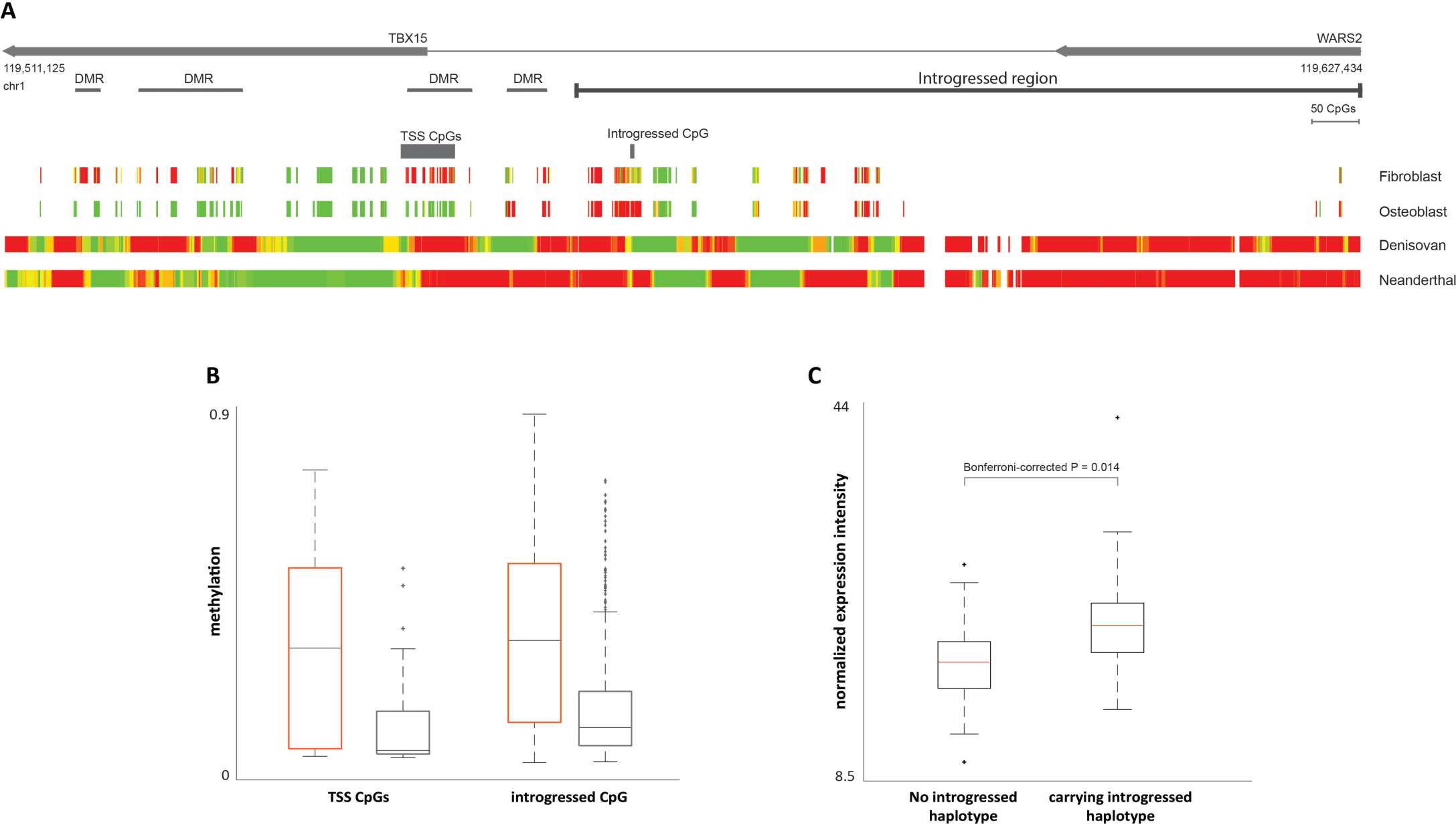
The regulatory effects of the introgressed haplotype on *TBX15* and *WARS2*. A. Methylation maps of the introgressed tract and its downstream region. The bottom panels show the reconstructed methylation maps of the Denisovan and the Neanderthal genomes, as well as two modern methylation maps from osteoblasts and fibroblasts. The maps are color coded from green (unmethylated) to red (methylated). The modern maps include fewer positions as they were produced using a reduced representation bisulfite sequencing (RRBS) protocol. Above the methylation maps are the CpG positions whose methylation levels are significantly associated with the introgressed haplotype. The top panels show *TBX15* and *WARS2*, the introgressed tract (as defined by the CRF), and the previously identified DMRs, where the Denisovan genome defers from present-day humans. B. Individuals carrying the introgressed haplotype (marked by black boxes) show lower levels of DNA methylation in both regions, compared to individuals who do not carry the introgressed haplotype (marked by orange boxes), in both the 16-CpG region around the TSS of *TBX15* (left), and a CpG position within the introgressed region (right). C. Expression of *TBX15* in individuals carrying the introgressed haplotype vs. individuals who do not carry it. *TBX15* is expressed on average at 22% higher levels in individuals carrying the haplotype.

To this end, we studied a data set of skin fibroblasts, where both genes are active, and in which each individual is characterized for sequence, methylation and expression. This data set included 62 individuals, of which two are homozygous for the introgressed tract, 21 are heterozygous, and 39 do not carry the tract (Wagner, et al. 2014). Due to the low number of individuals who are homozygous for the introgressed tract, we divided the individuals into two groups: individuals who do not carry the tract, and those who carry at least one copy of it. Then, we searched for CpG positions whose methylation level differs between the groups. Searching the entire region between the promoter of *WARS2* (5kb upstream to the TSS) to the transcription termination site of *TBX15*, we identified two such regions (FDR < 0.05, t-test, Table S3). The first region includes 16 CpGs around the TSS of the long transcript of *TBX15* (hereinafter “TSS CpGs”), all of which are significantly hypomethylated in introgressed individuals compared to individuals who do not carry the introgressed tract (Figure 7B). Interestingly, these CpGs almost completely overlap one of the previously reported DMRs (Figure 7A). The second region is a single CpG position that is found within the introgressed region, 11 kb upstream of the TSS of *TBX15* (hereinafter “upstream CpG”). Here too, introgressed individuals are hypomethylated compared to individuals who do not carry the tract (Figure 7B, Supplementary Table X).

In order to further test the effect of introgression on the activity of *TBX15* and *WARS2*, we investigated whether the methylation levels of the 16 TSS CpGs and the upstream CpG are associated with the levels of expression of *TBX15* and *WARS2*. We found that none of these CpGs are associated with the expression of *WARS2*, while all of them are significantly associated with the expression of *TBX15* (FDR < 0.05, Pearson correlation). In fact, both the TSS CpGs and the upstream CpG are found within two previously reported regulatory regions of *TBX15*, and it was shown that their methylation levels are linked to the activity levels of this gene (Chandra, et al. 2014). We also detect a more general association between the introgressed haplotype and expression levels: individuals carrying the haplotype exhibit a 22% increase in the expression level of *TBX15* in skin fibroblasts (Bonferroni-corrected P = 0.014, t-test, Figure 7C).

*TBX15* exhibits a unique link between expression and methylation, where, in contrast to most genes, global hypermethylation of its promoter is associated with elevated activity (Chandra, et al. 2014). However, the local relationship between the methylation at specific sites within its promoter and its activity level is substantially more complex, highly tissue-specific, and little understood to date. For example, unlike the rest of the promoter region, hypomethylation of the TSS region was shown to be associated with elevated expression levels of *TBX15* (Chandra, et al. 2014). This trend matches the one we see in the fibroblast data set, where introgressed individuals exhibit reduced methylation in the TSS CpGs and elevated expression levels of *TBX15* (Figure 7C).

A sensible assumption is that the methylation patterns in the introgressed individuals would resemble those in the Denisovan. However, the trend we observe is opposite; the Denisovan is hypermethylated compared to present-day humans, whereas introgressed individuals are hypomethylated compared to other present-day human individuals. While counterintuitive at first glance, it is important to take into account the complex and tissue-specific regulation of *TBX15*, and the fact that our analysis was conducted in fibroblasts, while the Denisovan methylation levels are from bone. Both the TSS CpGs and the upstream CpG reside in regions where methylation is variable across tissues (Chandra, et al. 2014). Moreover, the method of measurement of DNA methylation is different between these samples (high-resolution single site measurements in fibroblasts vs. regional reconstruction in the Denisovan bone). Therefore, further analyses are needed in order to determine how methylation patterns are affected by introgression in additional tissues, and specifically in bone.

The complex relationship between the introgressed tract and the activity levels of *TBX15* and *WARS2* can also be observed in the wider context of expression across tissues. To this end, we queried the GTEx database (GTEx-Consortium 2015) to examine if the introgressed SNPs are associated with an effect on human gene expression in various tissues. The sample sizes in the GTEx project do not provide enough power to detect trans-eQTLs, so we could only search for a cis-eQTL relationship with genes within a +/− 1 Mb window of each SNP. We therefore only checked whether the 28 introgressed SNPs that had signatures of positive selection in GI were also eQTLs for *TBX15* or *WARS2*. We found that all of these SNPs are tightly linked and therefore have almost exactly the same P-values in each of the tissues, so we only focus on one of these (rs2298080) below. Tables S4 and S5 show the effect sizes and P-values obtained from GTEx for 41 different tissues for *TBX15* and *WARS2*, respectively. In the case of *TBX15*, we find only one tissue with P < (0.05 / number of tissues tested), in the testis, where the Denisovan variant increases expression of the gene (P = 0.00018). When querying the SNP for expression effects on *WARS2*, we find 6 tissues with P < (0.05 / number of tissues tested): subcutaneous adipose, adrenal gland, aorta, tibial artery, esophagus muscularis, skeletal muscle. This suggests that expression differences, in the tissues and developmental stages represented in the GTEx database, are more ubiquitous in *WARS2* than *TBX15*. For all significant tissues, we find that the Denisovan variant decreases expression of *WARS2*. For *TBX15*, on the other hand, the trend varies between tissues.

### Association studies

Because the haplotype is at intermediate frequencies in Europeans, we queried the GIANT consortium GWAS data (Wood, et al. 2014; Locke, et al. 2015; Shungin, et al. 2015), which contains a number of anthropometric traits tested on a European panel. When looking at all P-values in the region, we find that there is a peak of significantly associated SNPs for three phenotypes right where the haplotype is inferred to be located (Figure S14). These phenotypes are BMI-adjusted waist circumference, waist-hip ratio and BMI-adjusted waist-hip ratio. We then queried the most extreme SNPs that serve to differentiate the archaic from the non-archaic haplotypes (see Methods) and the SNPs that were among the top PBS hits in GI (Table 1). The introgressed alleles in the queried SNPs are associated with a positive effect size for all three phenotypes. However, even though some of these SNPs are significantly associated with BMI-adjusted waist-circumference (P < 10^−5^), they are not among the top most significant SNPs in the region for any of the three phenotypes (Figure S14). Furthermore, when we conditioned on the SNP with the lowest P-value in the region (rs984222) we found no genome-wide significant association with the queried SNPs (lowest conditional P-value for introgressed SNPs = 0.047) (Figure S14). Interestingly, we also observed that the region overlaps a 100 kb region designated as a mouse QTL for the induction of brown adipocytes (MGI:2149993)(Xue, et al. 2005).

## Discussion

We have identified a highly divergent haplotype in the *TBX15/WARS2* region, which was likely introduced into the modern human gene pool via introgression with archaic humans. A priori, one would expect the source of introgression to be Neanderthals, due to their geographic distribution and the known admixture event(s) from Neanderthals into Eurasians (Green, et al. 2010; Prüfer, et al. 2014). However, the introgressed sequence is more closely related to the sequenced Denisovan genome than the sequenced Neanderthal genomes. This suggests that either the introgressing Neanderthal sequence is missing in the Neanderthals sequenced to date (and perhaps present in the sequenced Denisovan genome due to incomplete lineage sorting), or that the introgression event occurred from an unidentified population present in Eurasians that was more closely related to the Denisovan individuals found in the Altai Mountains than to any Neanderthal population.

The archaic haplotype is almost absent in Africans, present at higher frequencies in East Asians than in Europeans and South Asians, and at even higher frequencies in Native Americans and GI, where it is almost fixed (after correcting for post-Columbian admixture). This suggests there may have been a temporally and geographically extended period of selection for the archaic haplotype throughout eastern Eurasia. Population genetic differentiation values between Native Americans and Yoruba are significantly high (YRI vs. PEL: P < 10^−4^), but only marginally significant when comparing East Asians and Yoruba (YRI vs. CHB: P = 0.01), but only marginally significant when comparing East Asians and Native Americans (P = 0.02 for CHB vs. PEL and P = 0.06 for CHB vs. MXL). These findings suggest that selection on this locus may have acted during the early phases of the peopling of the Americas, perhaps in the Siberian or the Beringian ancestors of both modern Native Americans and Greenlandic Inuit. Selection may have continued to operate in Native Americans after their split from the Greenlandic Inuit, but we found that one need not invoke a model of continuing selection, if selection in Beringia was strong (2Ns > ~500) and/or the effective population size in Native Americans after their split was very small. While very strong selection is unlikely, effective population sizes for Native Americans have been estimated to be very low (Raghavan, et al. 2015).

It is intriguing that the introgressed tract is extremely long in 3 specific populations of central Asian and Siberia: the Yakut, the Even and the Naxi. The Yakut and Even are northeastern Siberian groups that migrated from the Lake Baikal and Transbaikal regions north of Mongolia. The Naxi are a population living in the Himalayan foothills that migrated from northwestern China. It is possible that these longer versions of the tract are perhaps remnants of the originally introgressed haplotype, which may have been shortened by recombination before sweeping to high frequencies in other populations. Though the haplotype is present in Europeans, there is some evidence to suggest it may have been introduced via the eastern steppe migrations of the Late Neolithic (Supplementary Information 4), again indicating that the original introgression event may have occurred somewhere in Asia.

The *TBX15/WARS2* region is highly pleiotropic: it has been found to be associated with a variety of traits. These include the differentiation of adipose tissue (Gburcik, et al. 2012), body fat distribution (Heid, et al. 2010; Liu, et al. 2013; Liu, et al. 2014; Shungin, et al. 2015), facial morphology (Lausch, et al. 2008; Pallares, et al. 2015), stature (Lausch, et al. 2008), ear morphology (Curry 1959; Adhikari, et al. 2015), hair pigmentation (Candille, et al. 2004) and skeletal development (Singh, et al. 2005; Lausch, et al. 2008). Interestingly, for several of the body fat distribution studies, the introgressed SNPs lie in the middle of a region with significant genome-wide associations, although the introgressed SNPs themselves do not have genome-wide significant P-values, after conditioning for the SNP with the strongest association in the region (which is not linked to the introgressed tract).

The haplotype is located immediately upstream of *TBX15*, overlapping some of its regulatory regions. Using fibroblast data where individuals were characterized for sequence, methylation and expression, we have shown a three-way association between the haplotype, the levels of DNA methylation, and the levels of expression of *TBX15*. However, the GTEx analysis found an association between the haplotype and the expression of *TBX15* only in the testis. Similarly, whereas we could not detect a regulatory link to the haplotype when analyzing *WARS2* in the fibroblast data, the GTEx analysis revealed that the Denisovan SNPs are cis-eQTLs for *WARS2* across various tissues. This included skeletal muscle and subcutaneous adipose. These contrasting results, together with the different methods used to perform measurements in these data sets, make it difficult to assess whether the downstream phenotypic changes are due to changes in the regulation of *TBX15, WARS2* or both.

Altogether, our study suggests a complex multi-factorial regulation of *TBX15* and *WARS2*. We show that the introgressed region is associated with regional changes in methylation and expression levels, but our findings also hint to other factors that affect the regulation of these genes that are yet to be elucidated.

## Methods

### f_D_ statistics in Greenlandic Inuit SNP data

Following refs. (Green, et al. 2010; Durand, et al. 2011), at site i, let Ci(ABBA) = ((1 − *f_Yoruba_*) × *f_Greenlandic Inuit_* × *f_Archaic human_*), where *f* is the derived allele frequency (with respect to the human-chimpanzee ancestor) in either a population panel (for the Greenlandic Inuit) or a diploid genome (for Yoruba and the archaic humans). Furthermore, let C_i_(BABA) = (*f_Yoruba_* × (1 − *f_Greenlandic Inuit_*) × *f_Archaic human_*). Then, for a set of N sites within a particular region of the genome, we computed D as follows:

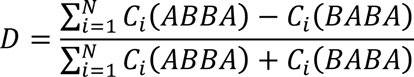

Let S(Yoruba,Greenlandic Inuit,Archaic,Chimpanzee) be the numerator in the D statistic defined above. We computed f_D_ as follows:

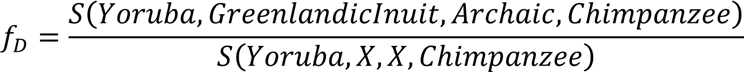

Here, X is defined – dynamically for each site i – as the population (either Archaic or Greenlandic Inuit) that has the highest derived allele frequency.

### Introgressed tracts in Eurasian whole-genome data

First, we used a HMM to find the archaic introgressed tracts in the region (Seguin-Orlando, et al. 2014)(Supplementary Information 1). We set an admixture proportion of 2%, an admixture time of 1900 generations ago and a constant recombination rate of 0.155 × 10^−8^ per bp per generation, which is the average recombination rate in the *TBX15/WARS2* region. We used YRI as the population with no introgression. The HMM was run individually on each phased genome from the 1000 Genomes data, using all SNPs that were variable in the continental panel to which the individual belonged. The model has two hidden states – introgressed and non-introgressed – and the rate of transition between the two assumes an exponential distribution of admixture tracts. We called tracts if the posterior probability for introgression estimated using the HMM was higher than 90%. We also tried increasing the admixture proportion to 10% and 50%, but did not observe any major differences in the length of the tracts or the proportion of individuals carrying them, except that some of the tracts in the same chromosome tended to be broken into smaller tracks more often than when using a 2% rate. Under these parameters, the HMM has a specificity of 99.56%, a sensitivity of 36.07% and a false discovery rate of 1.15%. We also obtained introgression tracks in the same region from Sriram Sankararaman (pers. comm.). These tracks were estimated using a CRF framework applied to the SGDP data(Sankararaman, et al. 2014; Sankararaman, et al. 2016). This method searches the genome for runs of archaic alleles that are of a length consistent with introgression.

### Uniquely shared sites

We defined “Eurasian uniquely shared sites” (Racimo, et al. 2016) as sites where the Denisovan genome is homozygous and where the Denisovan allele is at low frequency (< 1%) in Africans (AFR, excluding admixed African-Americans), but at high frequency (> 20%) in non-American Eurasians (EUR+EAS+SAS) from phase 3 of the 1000 Genomes Project (Auton, et al. 2015). Similarly, we defined the “derived shared quantile” statistic (Racimo, et al. 2016) as the 95% quantile of all derived allele frequencies in Eurasians, for SNPs where the Denisovan allele is homozygous for the derived allele and where the derived allele is at low frequency (< 1%) in Africans. In both cases, we only used sites that were not in repeat-masked regions (Smit, et al. 1996-2010.) and that lied in regions with 20-bp Duke mappability equal to 1 (Derrien, et al. 2012). The mapability track ensures that the sites are located in uniquely mappable 20-bp windows of the genome, to avoid issues that may spring from mismapped reads.

### Haplotype clustering

To examine the haplotypes in this region, we computed the number of pairwise differences between every pair of haplotypes in a particular continental panel. Then we ordered the haplotypes based on their number of pairwise distances to the archaic sequence in each continent. Figure 4 is generated using the heatmap.2 function from the gplots package of the statistical computing platform R (R-Core-Team 2012).

### Haplotype network

We built a haplotype network based on pair-wise differences using R (R-Core-Team 2012) and the software package pegas (Paradis 2010). To plot the network, we used the 20 most abundant present-day human haplotypes. To make a fair comparison with published distances for *EPAS1* (Huerta-Sanchez and Casey 2015), we looked at the 40 most abundant present-day haplotypes instead, and only counted differences at SNPs that were segregating in present-day humans. The network is produced using statistical parsimony, such that the most closely related haplotypes are connected first via the least number of mutations (“steps”)(Templeton, et al. 1992).

### F_ST_ scan in Native Americans

We extracted sequencing data in form of BAM for a 100 Mbp region surrounding the putative introgressed haplotype from the 1000 Genomes Project data set (Auton, et al. 2015). First, we selected all unrelated individuals belonging to 4 population panels: African Yoruba (YRI), Central Europeans from Utah (CEU), Han Chinese (CHB) and Peruvians from Lima (PEL). To select PEL individuals that would serve as optimal representatives of Native American genetic variation, we calculated admixture proportion among all YRI, CEU, CHB and PEL individuals assuming 4 ancestral populations using *NGSadmix* (Skotte, et al. 2013). We then extracted the first 30 PEL individuals showing the highest proportion of inferred Native American ancestry (> 89%). Similarly, we selected the first 30 individuals ranked by African, European and East Asian ancestry, respectively, and used each of these four 30-individual cohorts in the principal component analyses described below. For this analysis, we processed a total of 120 individuals. We then repeated the same procedure using Colombian (CLM), Mexican (MXL) or Puerto Rican (PUR) panels. After calculating the ancestry proportion as described above, and given the higher European admixture proportions for these populations, only 20 individuals for MXL, CLM or PUR were chosen as representatives of Native American variation (Native American ancestry component > 64%). To visually verify whether we correctly selected unadmixed individuals in the Latin American cohorts, we performed a principal component analysis using *ngsTools* (Fumagalli, et al. 2014). As only PEL and MXL selected individuals explained more than 70% of Native ancestry (Figure S6), we did not consider other Latin American populations for further analyses.

We computed *F_ST_*, a measure of population genetic differentiation, between the Latin Americans with the highest proportion of Native American ancestry and other populations against YRI, using a method-of-moments estimator implemented in *VCFtools* (Danecek, et al. 2011) from VCF files from the 1000 Genomes Project (Figure 5). To identify signatures of positive selection in Native Americans, we scanned the region around the putatively introgressed haplotype using a sliding-windows approach, with window size of 20kbp and step size of 2kbp, and computing *F_ST_* against YRI in each window. We also performed this same analysis but using all non-American, non-African populations present in the 1000 Genomes Project, by randomly sampling 30 individuals at each population for ease of comparison (Figure S7). To further investigate the deviation of the observed *F_ST_* from neutral expectations, we calculated *F_ST_* values for a marker for the introgressed haplotype (rs2298080) between all pairs of populations investigated for all reported biallelic SNPs across the whole genome (Table S4).

We were also interested in determining which demographic scenarios could generate levels of *F_ST_* between PEL and CHB as extreme as those observed, without invoking selection after the split between PEL and GI. We explored several combinations of effective population size for Native Americans (from 1,000 to 10,000 with a step size of 1,000) and split time between Native Americans and Greenlandic Inuit (from 8,000 to 18,000 years ago with a step size of 1,000 years). All other parameters were fixed to those inferred using *dadi* (Gutenkunst, et al. 2009). Additionally, we assumed that selection started in the ancestral population of Native Americans and Greenlandic Inuit 19,000 years ago. After the split, selection stopped in Native Americans, and lasted in the Greenlandic Inuit for 500 more years after the split. We tried a range of selection coefficients: 0, 20, 50, 200, 500, or 1,000 (in units of 2Ns). For each scenario, we simulated 10,000 replicates of a 20kbp region, using the software *msms (Ewing and Hermisson 2010)*, and recorded the 95th percentile of the *F_ST_* distribution between Native Americans and East Asians.

### TBX15 and WARS2 regulation

Data from SNP arrays, methylation arrays and expression arrays for the 62 fibroblast samples were downloaded from Gene Expression Omnibus (GEO accession number GSE53261). Expression values were normalized using Median Absolute Deviation (MAD) scale normalization (Fundel, et al. 2008). Chromatin peaks (H3K27ac, H3K4me1 and DNase-I) were downloaded from the Roadmap integrative analysis of 111 epigenomes (Kundaje, et al. 2015). For the analysis of the link between introgression and the expression level of TBX15, we corrected the t-test P-value using Bonferroni correction, taking into account the eight GTEx mesenchymal tissues in which TBX15 is expressed. Osteoblast and fibroblast RRBS maps were downloaded from the ENCODE project (GEO accession number: GSE27584).

### Conditional association study

We used the method of conditional association testing based on summary statistics (Yang, et al. 2012) that is implemented in the GCTA software package (Yang, et al. 2011). To obtain information about patterns of linkage disequilibrium for the European population analyzed by GIANT– which is required by the method – we used the called genotypes for the CEU panel from phase 3 of the 1000 Genomes Project.

## Acknowledgements

We thank Montgomery Slatkin and members of the Slatkin and Nielsen labs for helpful advice and discussions. We also thank Jacob Crawford for his help and advice in inferring admixture tracts using a hidden Markov model. Additionally, we thank Sriram Sankararaman for providing us with the output of his conditional random field method in our region of interest. RN is supported by a National Institutes of Health grant (R01HG003229). LC is supported by the Israel Science Foundation FIRST individual grant (ISF 1430/13). MF is supported by a Human Frontier Science Program fellowship (LT00320/2014). E.H.S is supported by UC Merced start-up funds and an NSF-DEB 1557151 grant.

## References

Abi-Rached L, Jobin MJ, Kulkarni S, McWhinnie A, Dalva K, Gragert L, Babrzadeh F, Gharizadeh B, Luo M, Plummer FA, et al. 2011. The shaping of modern human immune systems by multiregional admixture with archaic humans. Science 334:89–94.

Adhikari K, Reales G, Smith AJ, Konka E, Palmen J, Quinto-Sanchez M, Acuña-Alonzo V, Jaramillo C, Arias W, Fuentes M, et al. 2015. A genome-wide association study identifies multiple loci for variation in human ear morphology. Nat Commun 6:7500.

Allentoft ME, Sikora M, Sjögren KG, Rasmussen S, Rasmussen M, Stenderup J, Damgaard PB, Schroeder H, Ahlström T, Vinner L, et al. 2015. Population genomics of Bronze Age Eurasia. Nature 522:167–172.

Auton A, Brooks LD, Durbin RM, Garrison EP, Kang HM, Korbel JO, Marchini JL, McCarthy S, McVean GA, Abecasis GR, et al. 2015. A global reference for human genetic variation. Nature 526:68–74.

Candille SI, Van Raamsdonk CD, Chen C, Kuijper S, Chen-Tsai Y, Russ A, Meijlink F, Barsh GS. 2004. Dorsoventral patterning of the mouse coat by Tbx15. PLoS Biol 2:E3.

Chandra S, Baribault C, Lacey M, Ehrlich M. 2014. Myogenic differential methylation: diverse associations with chromatin structure. Biology (Basel) 3:426–451.

GTEx-Consortium. 2015. Human genomics. The Genotype-Tissue Expression (GTEx) pilot analysis: multitissue gene regulation in humans. Science 348:648–660.

Curry GA. 1959. Genetical and developmental studies on droopy-eared mice. Development 7:39–65.

Danecek P, Auton A, Abecasis G, Albers CA, Banks E, DePristo MA, Handsaker RE, Lunter G, Marth GT, Sherry ST, et al. 2011. The variant call format and VCFtools. Bioinformatics 27:2156–2158.

Dannemann M, Andrés AM, Kelso J. 2015. Adaptive variation in human toll-like receptors is contributed by introgression from both Neandertals and Denisovans. bioRxiv.

Derrien T, Estellé J, Marco Sola S, Knowles DG, Raineri E, Guigó R, Ribeca P. 2012. Fast computation and applications of genome mappability. PLoS One 7:e30377.

Diederichs S, Bartsch L, Berkmann JC, Fröse K, Heitmann J, Hoppe C, Iggena D, Jazmati D, Karschnia P, Linsenmeier M, et al. 2016. The dark matter of the cancer genome: aberrations in regulatory elements, untranslated regions, splice sites, non-coding RNA and synonymous mutations. EMBO Mol Med 8:442–457.

Durand EY, Patterson N, Reich D, Slatkin M. 2011. Testing for ancient admixture between closely related populations. Molecular biology and evolution 28:2239–2252.

Ewing G, Hermisson J. 2010. MSMS: a coalescent simulation program including recombination, demographic structure and selection at a single locus. Bioinformatics 26:2064–2065.

Fu Q, Li H, Moorjani P, Jay F, Slepchenko SM, Bondarev AA, Johnson PL, Aximu-Petri A, Prüfer K, de Filippo C, et al. 2014. Genome sequence of a 45,000-year-old modern human from western Siberia. Nature 514:445–449.

Fumagalli M, Moltke I, Grarup N, Racimo F, Bjerregaard P, Jørgensen ME, Korneliussen TS, Gerbault P, Skotte L, Linneberg A, et al. 2015. Greenlandic Inuit show genetic signatures of diet and climate adaptation. Science 349:1343–1347.

Fumagalli M, Vieira FG, Linderoth T, Nielsen R. 2014. ngsTools: methods for population genetics analyses from next-generation sequencing data. Bioinformatics 30:1486–1487.

Fundel K, Haag J, Gebhard PM, Zimmer R, Aigner T. 2008. Normalization strategies for mRNA expression data in cartilage research. Osteoarthritis Cartilage 16:947–955.

Gamba C, Jones ER, Teasdale MD, McLaughlin RL, Gonzalez-Fortes G, Mattiangeli V, Dombróczki L, Kővári I, Pap I, Anders A, et al. 2014. Genome flux and stasis in a five millennium transect of European prehistory. Nat Commun 5:5257.

Gburcik V, Cawthorn WP, Nedergaard J, Timmons JA, Cannon B. 2012. An essential role for Tbx15 in the differentiation of brown and “brite” but not white adipocytes. Am J Physiol Endocrinol Metab 303:E1053–1060.

Gokhman D, Lavi E, Prüfer K, Fraga MF, Riancho JA, Kelso J, Pääbo S, Meshorer E, Carmel L. 2014. Reconstructing the DNA methylation maps of the Neandertal and the Denisovan. Science 344:523–527.

Gravel S. 2012. Population genetics models of local ancestry. Genetics 191:607–619.

Gravel S, Henn BM, Gutenkunst RN, Indap AR, Marth GT, Clark AG, Yu F, Gibbs RA, Bustamante CD, Project G. 2011. Demographic history and rare allele sharing among human populations. Proc Natl Acad Sci U S A 108:11983–11988.

Green RE, Krause J, Briggs AW, Maricic T, Stenzel U, Kircher M, Patterson N, Li H, Zhai W, Fritz MH, et al. 2010. A draft sequence of the Neandertal genome. Science 328:710–722.

Gutenkunst RN, Hernandez RD, Williamson SH, Bustamante CD. 2009. Inferring the joint demographic history of multiple populations from multidimensional SNP frequency data. PLoS Genet 5:e1000695.

Haak W, Lazaridis I, Patterson N, Rohland N, Mallick S, Llamas B, Brandt G, Nordenfelt S, Harney E, Stewardson K, et al. 2015. Massive migration from the steppe was a source for Indo-European languages in Europe. Nature 522:207–211.

Heid IM, Jackson AU, Randall JC, Winkler TW, Qi L, Steinthorsdottir V, Thorleifsson G, Zillikens MC, Speliotes EK, Mägi R, et al. 2010. Meta-analysis identifies 13 new loci associated with waist-hip ratio and reveals sexual dimorphism in the genetic basis of fat distribution. Nat Genet 42:949–960.

Hellenthal G, Stephens M. 2007. msHOT: modifying Hudson’s ms simulator to incorporate crossover and gene conversion hotspots. Bioinformatics 23:520–521.

Huerta-Sanchez E, Casey FP. 2015. Archaic inheritance: supporting high altitude life in Tibet. Journal of Applied Physiology 119:1129–1134.

Huerta-Sánchez E, Jin X, Asan, Bianba Z, Peter BM, Vinckenbosch N, Liang Y, Yi X, He M, Somel M, et al. 2014. Altitude adaptation in Tibetans caused by introgression of Denisovan-like DNA. Nature 512:194–197.

International-HapMap-Consortium. 2005. A haplotype map of the human genome. Nature 437:1299–1320.

Korneliussen TS, Albrechtsen A, Nielsen R. 2014. ANGSD: Analysis of Next Generation Sequencing Data. BMC Bioinformatics 15:356.

Kron K, Liu L, Trudel D, Pethe V, Trachtenberg J, Fleshner N, Bapat B, van der Kwast T. 2012. Correlation of ERG expression and DNA methylation biomarkers with adverse clinicopathologic features of prostate cancer. Clin Cancer Res 18:2896–2904.

Kundaje A, Meuleman W, Ernst J, Bilenky M, Yen A, Heravi-Moussavi A, Kheradpour P, Zhang Z, Wang J, Ziller MJ, et al. 2015. Integrative analysis of 111 reference human epigenomes. Nature 518:317–330.

Lausch E, Hermanns P, Farin HF, Alanay Y, Unger S, Nikkel S, Steinwender C, Scherer G, Spranger J, Zabel B, et al. 2008. TBX15 mutations cause craniofacial dysmorphism, hypoplasia of scapula and pelvis, and short stature in Cousin syndrome. Am J Hum Genet 83:649–655.

Lazaridis I, Patterson N, Mittnik A, Renaud G, Mallick S, Kirsanow K, Sudmant PH, Schraiber JG, Castellano S, Lipson M, et al. 2014. Ancient human genomes suggest three ancestral populations for present-day Europeans. Nature 513:409–413.

Liang M, Nielsen R. 2014. The Lengths of Admixture Tracts. Genetics 197:953–967.

Liu CT, Buchkovich ML, Winkler TW, Heid IM, Borecki IB, Fox CS, Mohlke KL, North KE, Adrienne Cupples L, Consortium AAAG, et al. 2014. Multi-ethnic fine-mapping of 14 central adiposity loci. Hum Mol Genet 23:4738–4744.

Liu CT, Monda KL, Taylor KC, Lange L, Demerath EW, Palmas W, Wojczynski MK, Ellis JC, Vitolins MZ, Liu S, et al. 2013. Genome-wide association of body fat distribution in African ancestry populations suggests new loci. PLoS Genet 9:e1003681.

Locke AE, Kahali B, Berndt SI, Justice AE, Pers TH, Day FR, Powell C, Vedantam S, Buchkovich ML, Yang J, et al. 2015. Genetic studies of body mass index yield new insights for obesity biology. Nature 518:197–206.

Mansour MR, Abraham BJ, Anders L, Berezovskaya A, Gutierrez A, Durbin AD, Etchin J, Lawton L, Sallan SE, Silverman LB, et al. 2014. Oncogene regulation. An oncogenic super-enhancer formed through somatic mutation of a noncoding intergenic element. Science 346:1373–1377.

Marcus JH, Novembre J. 2016. Visualizing the Geography of Genetic Variants. bioRxiv.

Martin SH, Davey JW, Jiggins CD. 2014. Evaluating the use of ABBA–BABA statistics to locate introgressed loci. Molecular biology and evolution.

Messer PW. 2013. SLiM: simulating evolution with selection and linkage. Genetics 194:1037–1039.

Meyer M, Kircher M, Gansauge MT, Li H, Racimo F, Mallick S, Schraiber JG, Jay F, Prüfer K, de Filippo C, et al. 2012. A high-coverage genome sequence from an archaic Denisovan individual. Science 338:222–226.

Moltke I, Fumagalli M, Korneliussen TS, Crawford JE, Bjerregaard P, Jørgensen ME, Grarup N, Gulløv HC, Linneberg A, Pedersen O, et al. 2015. Uncovering the genetic history of the present-day Greenlandic population. Am J Hum Genet 96:54–69.

Pallares LF, Carbonetto P, Gopalakrishnan S, Parker CC, Ackert-Bicknell CL, Palmer AA, Tautz D. 2015. Mapping of Craniofacial Traits in Outbred Mice Identifies Major Developmental Genes Involved in Shape Determination. PLoS Genet 11:e1005607.

Paradis E. 2010. pegas: an R package for population genetics with an integrated-modular approach. Bioinformatics 26:419–420.

Paten B, Herrero J, Beal K, Fitzgerald S, Birney E. 2008. Enredo and Pecan: genome-wide mammalian consistency-based multiple alignment with paralogs. Genome Res 18:1814–1828.

Prüfer K, Racimo F, Patterson N, Jay F, Sankararaman S, Sawyer S, Heinze A, Renaud G, Sudmant PH, de Filippo C, et al. 2014. The complete genome sequence of a Neanderthal from the Altai Mountains. Nature 505:43–49.

Racimo F, Sankararaman S, Nielsen R, Huerta-Sánchez E. 2015. Evidence for archaic adaptive introgression in humans. Nat Rev Genet 16:359–371.

Raghavan M, Skoglund P, Graf KE, Metspalu M, Albrechtsen A, Moltke I, Rasmussen S, Stafford TW, Orlando L, Metspalu E, et al. 2014. Upper Palaeolithic Siberian genome reveals dual ancestry of Native Americans. Nature 505:87–91.

Raghavan M, Steinrücken M, Harris K, Schiffels S, Rasmussen S, DeGiorgio M, Albrechtsen A, Valdiosera C, Ávila-Arcos MC, Malaspinas AS, et al. 2015. Genomic evidence for the Pleistocene and recent population history of Native Americans. Science 349:aab3884.

R-Core-Team. 2012. R: A Language and Environment for Statistical Computing. In. Vienna, Austria.

Sankararaman S, Mallick S, Dannemann M, Prüfer K, Kelso J, Pääbo S, Patterson N, Reich D. 2014. The genomic landscape of Neanderthal ancestry in present-day humans. Nature 507:354–357.

Sankararaman S, Mallick S, Patterson N, Reich D. 2016. The Combined Landscape of Denisovan and Neanderthal Ancestry in Present-Day Humans. Curr Biol 26:1241–1247.

Seguin-Orlando A, Korneliussen TS, Sikora M, Malaspinas AS, Manica A, Moltke I, Albrechtsen A, Ko A, Margaryan A, Moiseyev V, et al. 2014. Genomic structure in Europeans dating back at least 36,200 years. Science.

Shungin D, Winkler TW, Croteau-Chonka DC, Ferreira T, Locke AE, Mägi R, Strawbridge RJ, Pers TH, Fischer K, Justice AE, et al. 2015. New genetic loci link adipose and insulin biology to body fat distribution. Nature 518:187–196.

Singh MK, Petry M, Haenig B, Lescher B, Leitges M, Kispert A. 2005. The T-box transcription factor Tbx15 is required for skeletal development. Mech Dev 122:131–144.

Skotte L, Korneliussen TS, Albrechtsen A. 2013. Estimating individual admixture proportions from next generation sequencing data. Genetics 195:693–702. *RepeatMasker Open-3.0* [Internet]. http://www.repeatmasker.org1996-2010.

Steegmann AT, Cerny FJ, Holliday TW. 2002. Neandertal cold adaptation: physiological and energetic factors. Am J Hum Biol 14:566–583.

Templeton AR, Crandall KA, Sing CF. 1992. A cladistic analysis of phenotypic associations with haplotypes inferred from restriction endonuclease mapping and DNA sequence data. III. Cladogram estimation. Genetics 132:619–633.

Vernot B, Akey JM. 2014. Resurrecting surviving Neandertal lineages from modern human genomes. Science 343:1017–1021.

Wagner JR, Busche S, Ge B, Kwan T, Pastinen T, Blanchette M. 2014. The relationship between DNA methylation, genetic and expression inter-individual variation in untransformed human fibroblasts. Genome Biol 15:R37.

Wood AR, Esko T, Yang J, Vedantam S, Pers TH, Gustafsson S, Chu AY, Estrada K, Luan J, Kutalik Z, et al. 2014. Defining the role of common variation in the genomic and biological architecture of adult human height. Nat Genet 46:1173–1186.

Xue B, Coulter A, Rim JS, Koza RA, Kozak LP. 2005. Transcriptional synergy and the regulation of Ucp1 during brown adipocyte induction in white fat depots. Mol Cell Biol 25:8311–8322.

Yang J, Ferreira T, Morris AP, Medland SE, Madden PA, Heath AC, Martin NG, Montgomery GW, Weedon MN, Loos RJ, et al. 2012. Conditional and joint multiple-SNP analysis of GWAS summary statistics identifies additional variants influencing complex traits. Nat Genet 44:369-375, S361–363.

Yang J, Lee SH, Goddard ME, Visscher PM. 2011. GCTA: a tool for genome-wide complex trait analysis. Am J Hum Genet 88:76–82.

Yi X, Liang Y, Huerta-Sanchez E, Jin X, Cuo ZX, Pool JE, Xu X, Jiang H, Vinckenbosch N, Korneliussen TS, et al. 2010. Sequencing of 50 human exomes reveals adaptation to high altitude. Science 329:75–78.

